# Genome scale metabolic network modelling for metabolic profile predictions

**DOI:** 10.1101/2023.07.26.550622

**Authors:** Juliette Cooke, Maxime Delmas, Cecilia Wieder, Pablo Rodriguez Mier, Clément Frainay, Florence Vinson, Timothy Ebbels, Nathalie Poupin, Fabien Jourdan

**Affiliations:** Toxalim (Research Centre in Food Toxicology), Université de Toulouse, INRAE, ENVT, INP-Purpan, UPS, Toulouse, France; Idiap Research Institute, Martigny, Switzerland; Section of Bioinformatics, Division of Systems Medicine, Department of Metabolism, Digestion, and Reproduction, Faculty of Medicine, Imperial College London, London, United Kingdom; Heidelberg University, Faculty of Medicine, Heidelberg University Hospital, Institute for Computational Biomedicine, Bioquant, Heidelberg, Germany; MetaToul-MetaboHUB, National Infrastructure of Metabolomics and Fluxomics, Toulouse, France

## Abstract

Metabolic profiling (metabolomics) aims at measuring small molecules (metabolites) in complex samples like blood or urine for human health studies. While biomarker-based assessment often relies on a single molecule, metabolic profiling combines several metabolites to create a more complex and more specific fingerprint of the disease. However, in contrast to genomics, there is no unique metabolomics setup able to measure the entire metabolome. This challenge leads to tedious and resource consuming preliminary studies to be able to design the right metabolomics experiment. In that context, computer assisted metabolic profiling can be of strong added value to design more quickly and efficiently metabolomics studies. We propose a constraint-based modelling approach which predicts *in silico* profiles of metabolites that are more likely to be differentially abundant under a given metabolic perturbation (*e.g.* due to a genetic disease), using flux simulation. In genome-scale metabolic networks, the fluxes of exchange reactions, also known as the flow of metabolites through their external transport reactions, can be simulated and compared between control and disease conditions in order to calculate changes in metabolite import and export. These import/export flux differences would be expected to induce changes in circulating biofluid levels of those metabolites, which can then be interpreted as potential biomarkers or metabolites of interest. In this study, we present SAMBA (SAMpling Biomarker Analysis), an approach which simulates fluxes in exchange reactions following a metabolic perturbation using random sampling, compares the simulated flux distributions between the baseline and modulated conditions, and ranks predicted differentially exchanged metabolites as potential biomarkers for the perturbation. We show that there is a good fit between simulated metabolic exchange profiles and experimental differential metabolites detected in plasma, such as patient data from the disease database OMIM, and metabolic trait-SNP associations found in mGWAS studies. These biomarker recommendations can provide insight into the underlying mechanism or metabolic pathway perturbation lying behind observed metabolite differential abundances, and suggest new metabolites as potential avenues for further experimental analyses.

**Author summary:** Associating diseases and other metabolic disruptions with physiological markers is key for diagnostic and personalised medicine. These markers can be metabolites - small molecules involved in every living being’s metabolism, and can be measured in biofluids such as blood or urine using metabolic profiling (metabolomics). Nevertheless, this experimental metabolomics design needs to be tailor made for each disease to ensure that most relevant metabolites will be detected. The selection of metabolites to analyse for future experiments can be time-consuming and expensive. In this paper, we build upon an existing computational method for simulating metabolite changes in a human model. This provides a prediction of the change in biofluid abundance of every known metabolite involved in human metabolism in a potentially large number of metabolic situations. The newly introduced method produces a change score and a rank for each metabolite in each condition. We show the strong potential of the approach by comparing predictions with experimental results.

## Introduction

Molecular biomarkers are measurable molecules directly or indirectly related to alterations of a certain physiological state. They can be used as diagnostic indicators, and can be measured in order to detect the presence or severity of many different diseases [1]. Among them, metabolic biomarkers are small molecules (metabolites) whose concentrations differ from a healthy state due to changes in the organism’s or tissue’s metabolism. In clinical settings, these biomarkers are traditionally detected using targeted bioassays which result in measurements for a small number of well characterised diagnostic metabolites (Figure 1A), such as glucose for diabetic disorders.

**Figure 1:**
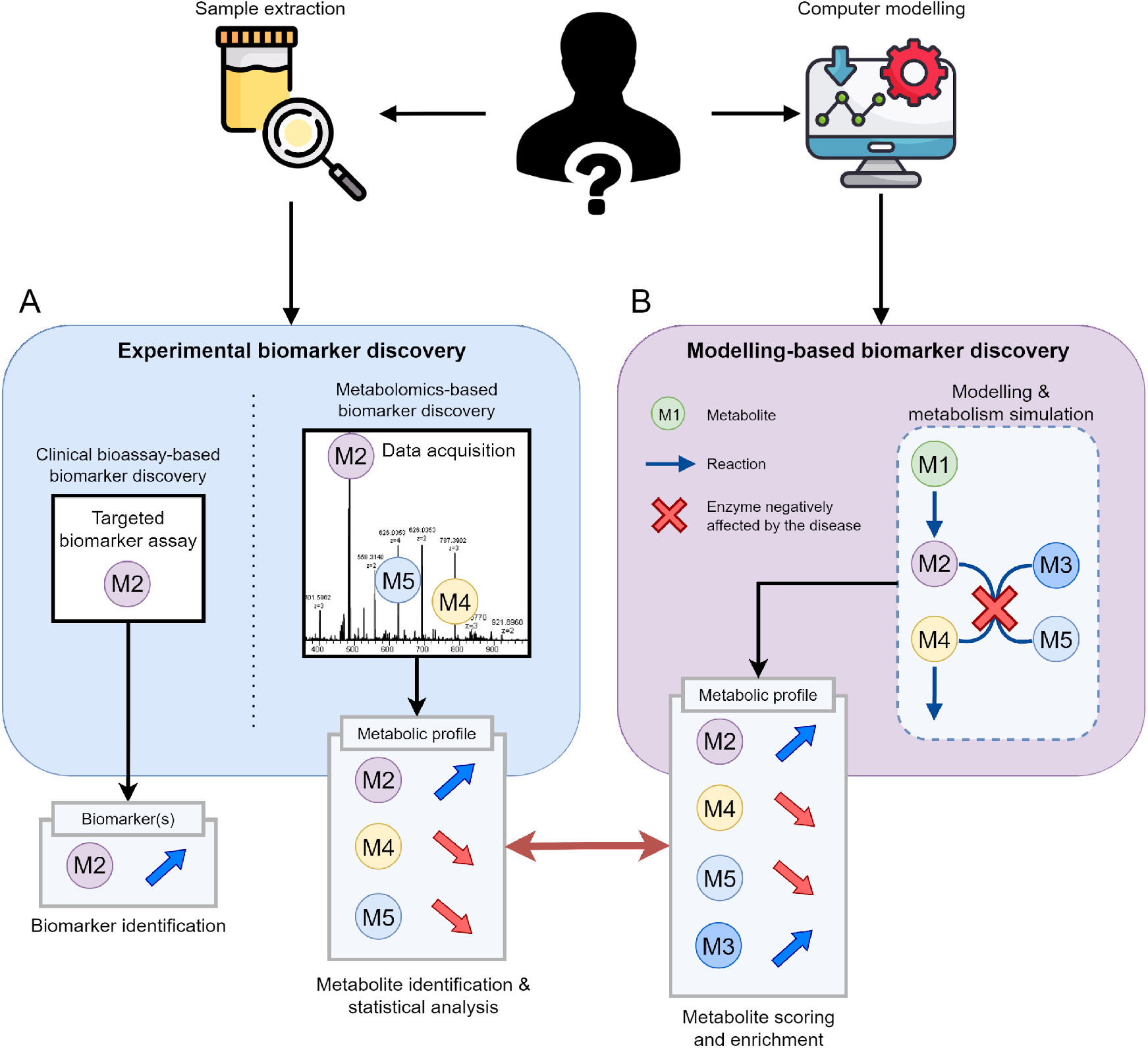
Biomarker discovery strategies, from targeted assays to in silico predictions. A: Experimental-based biomarker discovery produces targeted biomarkers or metabolic profiles. B: Metabolic disruptions can be modelled to simulate metabolic profiles similar to those generated using metabolomics.

A more holistic approach to measuring metabolites experimentally is via metabolic profiling, also called metabolomics [2]. In this paradigm, the detection of metabolic modulations relies on a broader panel of molecules (the metabolic profile) than targeted biochemical assays, hence resulting in an increased predictive and explanatory potential (Figure 1A). The most common analytical techniques used to measure metabolites are NMR (Nuclear Magnetic Resonance) and MS (Mass Spectrometry) coupled with separation techniques such as Liquid or Gas Chromatography.

Both NMR and MS are unable to completely cover the entire metabolome (see [3] for metabolic coverage assessment of MS data). In fact, both analytical setups have the ability to detect a portion of the metabolome depending on the physico-chemical properties of molecules present in the sample (*e.g.* polarity) [4]. Hence, it is important to select the right analytical setup, or combination of setups [5], to ensure that measured metabolites will be relevant for a specific disease. This experimental design step can be time and resource consuming for experts.

Even if a technique is in principle able to measure a relevant metabolite, it still requires an identification step to ensure the measured feature is a metabolite of interest. In MS for instance, the exact mass is not sufficient to confidently identify a metabolite. It requires confronting measurements with at least an orthogonal approach and confirming the identification by comparing the observed spectra with the one of the pure molecule (reference spectra of a standard, level 1 identification [6]). However, the reference spectra of the metabolite might not be available, which requires the laboratory to buy and inject new standards into mass spectrometers. Computational solutions can be used to fill the gaps in experimental observations, and facilitating the choice of standard molecules to measure using *in silico* methods would strongly benefit metabolomics laboratories.

A well suited class of methods for global metabolic modelling is Constraint-Based Modelling (CBM). CBM is a modelling framework which uses genome-scale metabolic networks, under the formalism of a stoichiometric matrix, to compute steady-state metabolic fluxes (the flow of metabolites) through biochemical reactions [7]. These networks aim to encompass all known metabolic genes, reactions and metabolites as well as the interactions between them for a given organism [8–10]. They can also be built to model a specific tissue or cell type [11], or even multiple tissues linked together [12].

CBM can be used to predict fluxes at steady-state under various conditions. This is achieved by defining metabolism as a system of linear mass balance equations, composed of the reaction flux vectors for each metabolite. These fluxes exist under defined flux constraints (setting upper and lower bounds) to model different metabolic states and reaction directionality. Controlling these bounds can for instance be used to simulate the complete knock-out (KO) (Figure 1B) of one or multiple gene(s) or reaction(s) (like for genetic diseases), or the reduction of the flux through a reaction or multiple reactions (knock-down), representing reduced enzyme activity due to some effect of treatment, regulation, or exposure to xenobiotics.

By using an organism-specific metabolic network in conjunction with a metabolic disruption, this CBM methodology can be used to predict which metabolites will be more or less released in biofluids. Indeed, in metabolic networks, some metabolites can be transported in and out from the internal compartment (cell or tissue) to the external compartment (e.g. biofluid or cell culture medium) usually using a single specific exchange reaction. For the *in silico* prediction of biomarkers these exchange reactions can be used to model the in/out flux of metabolites between tissues and circulating biofluids like blood or urine. This is why, in the context of metabolic profile prediction, the focus must be on these specific exchange reactions from the metabolic network in order to predict the equivalent of “biofluid metabolite level changes” using CBM. A break-down of this methodology is shown in Figure 2, using a simple metabolic network to compare flux simulations in healthy and disease conditions, and resulting in a ranked list of metabolites which change the most between the two conditions.

**Figure 2:**
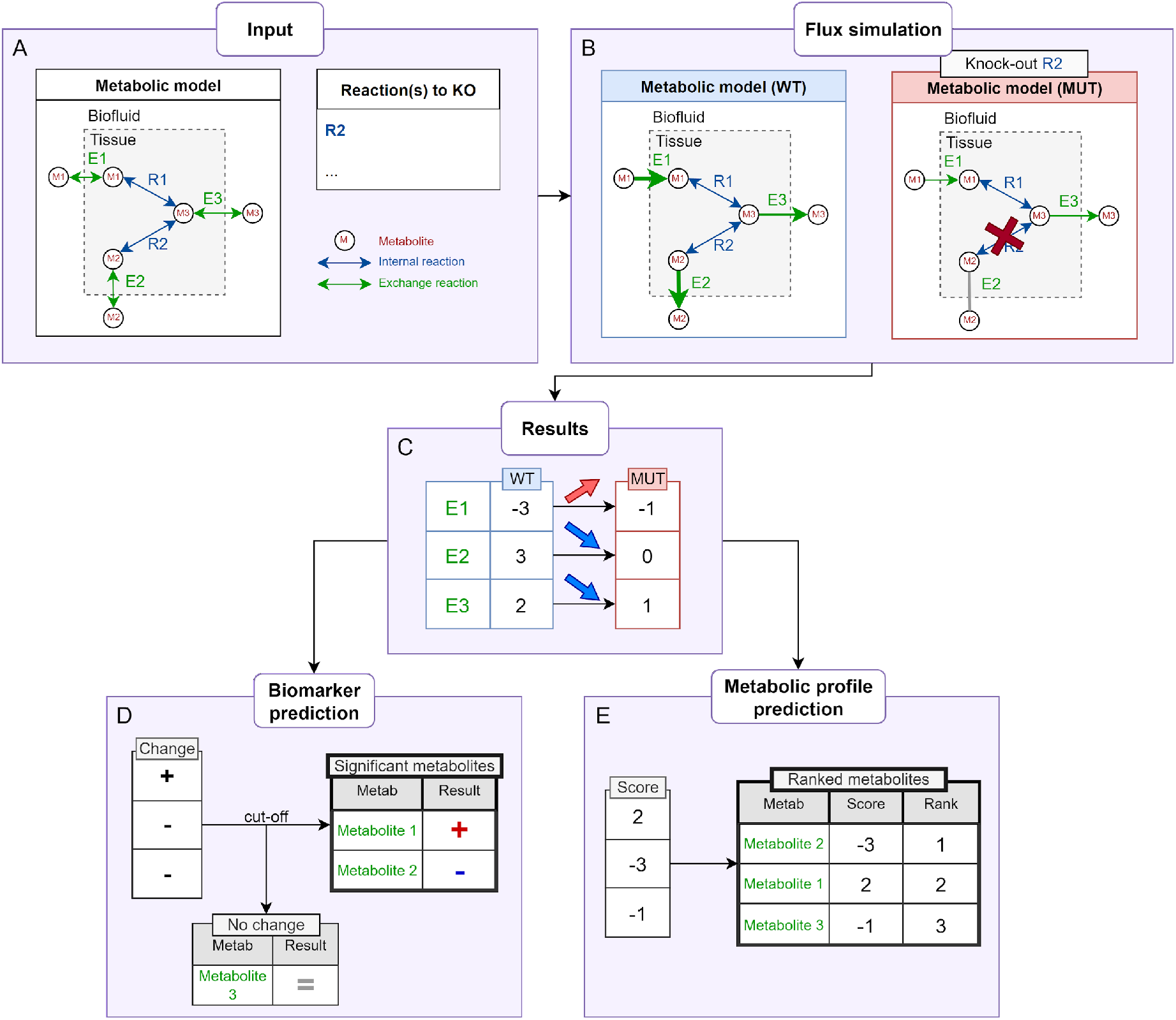
Methodology for the comparison of flux values and prediction of metabolite ranks using a simple network (A) in two conditions (B), with single flux values (C). The methodology from Shlomi *et al.* is shown in (D): each exported metabolite will have an associated change score for a given pair of conditions. (E) shows our methodology of scoring and ranking by absolute value among all of the metabolites in the network.

The overall methodology introduced in Shlomi *et al.* [13] for biomarker prediction consists in comparing the flux of exchange reactions between two conditions: one corresponding to a standard metabolic state (wild type) and one to a disease state. The wild type (WT) model is obtained by running the flux simulation on the chosen genome-scale metabolic network with default constraints, forcing the reaction of interest to have a non-zero flux (see Methods for more details). Then, the disease or mutant state model (MUT) is obtained by simulating a knock-out or knock down state using a list of genes or reactions to constrain (e.g. imposing null flux for knocked out reactions). Flux simulation is then also run on this MUT model (Figure 2B). In both states, fluxes are assessed simultaneously for all of the reactions in the network, however in the case of metabolic profile prediction, we only consider the exchange reaction fluxes. In constraint-based metabolic models a positive exchange reaction flux value represents an export of the corresponding metabolite, and a negative flux value means that the metabolite is imported. These flux values are represented in Figure 2B as the thickness of the arrows, and in Figure 2C as single flux values for each metabolite’s exchange reaction. Then, the flux values for every exchange reaction in the network can be compared between the WT and MUT conditions (Figure 2C) to determine a change (Figure 2D). The change of a given metabolite’s exchange reaction flux provides information on the corresponding metabolite’s production/consumption in both conditions, which leads over time to an increase, decrease or unchanged concentration in the biofluids. This is based on the assumption that the metabolites are not being depleted or produced through alternate external metabolic processes, or, at the very least, the rate of depletion/production is negligible enough to have any noticeable effect on the extracellular metabolic concentrations.

In Shlomi *et al.* and Thiele *et al.* [13,14], the authors show that CBM modelling can be used to predict biomarkers for diseases associated with gene deletions, validating these predictions with data from targeted clinical assays (focusing on specific classes of compounds like amino acids). The resulting predictions are boolean values associated with metabolites (*i.e.,* the metabolite is a biomarker or is not a biomarker (Figure 2D)), which does not allow the assessment of expected levels of changes in terms of concentrations.

In this article, we propose to go beyond previous work by predicting metabolic changes for 1497 metabolites from the Human 1 metabolic network. The method can also be used to pinpoint metabolites not detected in the selected analytical setup, therefore guiding metabolomics researchers in the selection and design of assays. To do so, we implement a measure which ranks metabolites based on their likelihood to be modulated under a given genetic, environmental, or otherwise metabolic stress (Figure 2E). To highlight the strong potential of our approach, we demonstrate how predicted metabolite rankings fit data from human metabolic profiling studies (both targeted and untargeted), hence paving the way for combining both *in silico* and experimental metabolic profiling.

This recommendation system is implemented in a freely available computational workflow at the following address: https://forgemia.inra.fr/metexplore/cbm/samba-project/samba.

## Results

### Sampling based flux estimation for improved metabolite change predictions

Metabolic network models are usually undetermined, *i.e.,* there are not enough constraints in the model to determine a unique solution for the mass balance system of linear equations [7]. For example, in the WT (Figure 2B), any other value for E1 would still ensure the steady state, as long as the other reactions’ flux values are changed accordingly to compensate. This is why it is difficult to define any one exact value for a reaction, and it is generally necessary to evaluate the range of possible flux values for all reactions through various CBM methods.

Figure 3 highlights two CBM methods for assessing the variability of flux values, again in two different conditions (WT and MUT). Flux Variability Analysis (FVA) [15] predicts minimum and maximum flux bounds for each reaction in the network (Figure 3B) under the predefined constraints. This interval represents the range of possible values that the fluxes of a given reaction can take. In an extreme case, like a complete KO or a single flux value, both the upper and lower bounds of the involved reaction will have null or close to zero values, like R2 in the MUT state in Figure 3B.

**Figure 3:**
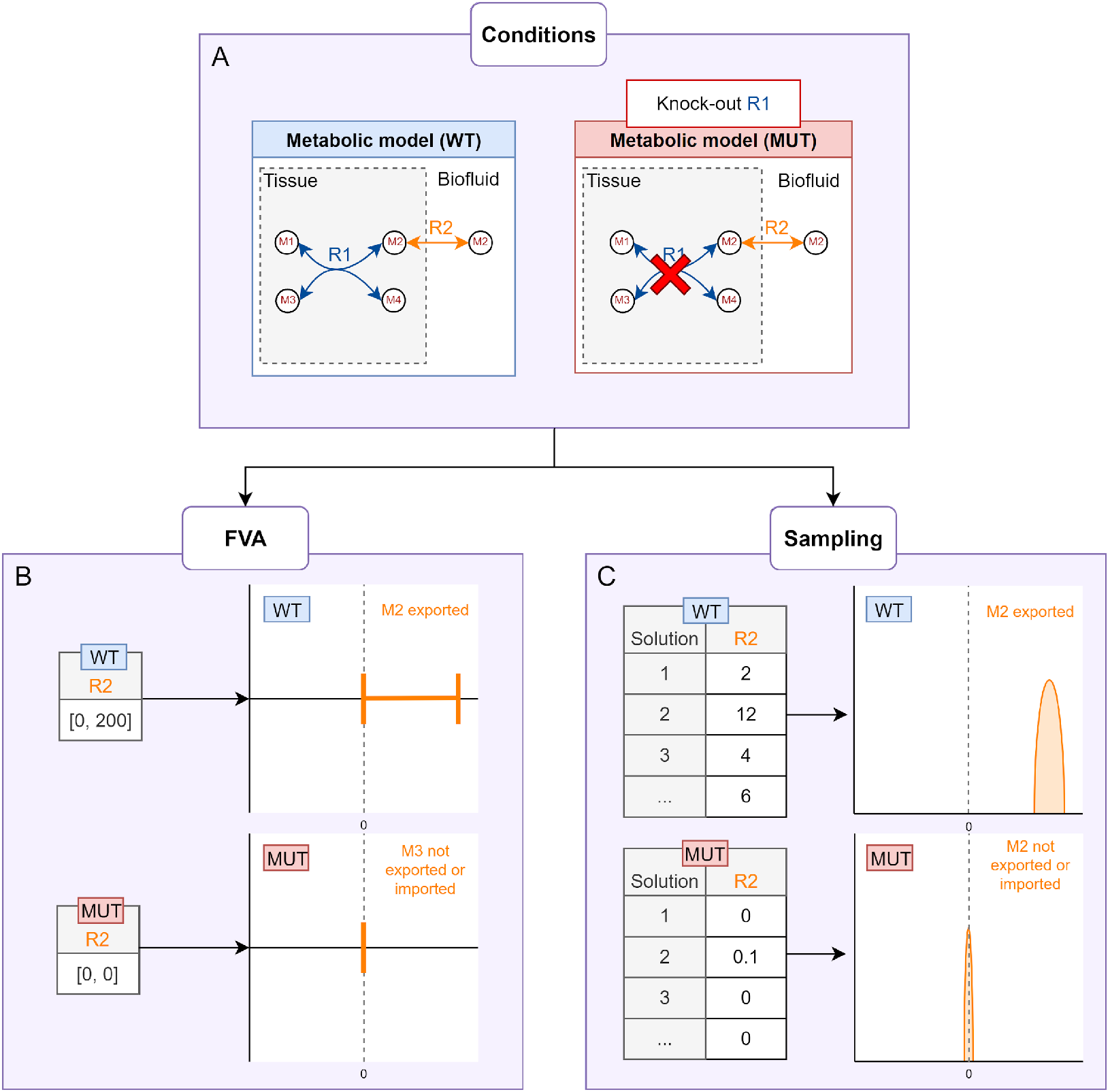
Flux Variability Analysis (FVA) and sampling for simulating fluxes in different conditions (A). The resulting flux values to be compared differ depending on the method used. FVA generates minimum and maximum possible flux values, shown as intervals (B), whereas sampling generates many values within those bounds, shown as distributions (C).

In Shlomi *et al.* and Thiele *et al.* [13,14], the authors used FVA to predict biomarkers by comparing two conditions: WT, and inborn errors of metabolism as gene knock-outs. If the FVA intervals of an exchange reaction shift between the two conditions, then the corresponding metabolite is considered to be a potential biomarker for the disease state (like for R2 in Figure 3B).

A second method for determining the most frequent flux values for a given reaction is by assessing the possible flux configurations (distributions) which fit the constraints defined in the model. These constraints on reaction fluxes (minimum and maximum possible values) define boundaries in the space of all possible flux values: the solution space. The solution space contains an infinite number of feasible flux values, hence it is impossible to fully explore the entire solution space without an infinite number of samples. Markov Chain Monte Carlo methods can be utilised for uniformly sampling the constrained space of feasible solutions, thereby offering a more comprehensive characterization of the solution space [16–18].

FVA may conceal the fact that the values present in most of the solution combinations (*e.g.* the most frequent flux values) are, for instance, close to one of the bounds as opposed to at the halfway point. For example, by comparing Figure 3B and C, the most frequent flux values for WT are much closer to the upper FVA bound than the centre of the bounds. It is also possible for two reactions to have the same extreme bounds but with flux values which are spread out differently in between the bounds. In other words, FVA is not sufficiently precise to assess the most frequent flux values. FVA does not provide insight into the distribution of flux values between those bounds since the most frequent flux value for a reaction could be located anywhere in between the two bounds.

By exploring the many combinations of flux values that satisfy the model constraints, sampling provides an overview of the most frequently valid flux values for every reaction in the network. Indeed, any one possible solution is a specific combination of flux values for each reaction, and some flux values are more frequent than others, meaning that they appear more often in different valid solutions. These many solutions can therefore outline each reaction’s flux distribution and thus reveal the most frequent fluxes along with the variability of the values, as shown in Figure 3C.

In this study, we propose an application we developed specifically for predicting metabolic profiles using random sampling, called SAMBA (SAMpling Biomarker Analysis) (using the optGPSampler algorithm).

### Scoring and ranking metabolites according to their predicted out/influx changes

While previous modelling methods had been used to predict biomarkers, the goal here is to predict entire metabolic profiles by capturing, for each metabolite, its amplitude of variation between the control and the condition under study. This variability can be evaluated, and metabolites can be scored and ranked in order to prioritise the ones to be measured and annotated during metabolic profiling.

SAMBA is used to sample flux distributions for every exchange reaction in the network, in two network states (WT and MUT). A score is calculated for each exchange reaction in the model by comparing the samples between both states. We propose the use of a z-score to evaluate the shift in distributions weighted by their variance, based on the z-score used in Mo *et al.* [19]. Z-scores are calculated for each metabolite’s exchange reaction ex_i_ between the WT and MUT. First, we sample a number (by default the total number of samples) of random pairs of values from the WT and MUT distributions. The collections of all MUT samples and WT samples for metabolite i’s exchange reaction are MUT_i_ and WT_i_ respectively. A “difference distribution” dd_i_ is calculated by subtracting random pairs of values from both MUT_i_ and WT_i_ (Equation 1). These random samples are not matched: the two WT and MUT values are not necessarily from the same sample step. The final z-score z_i_ is calculated by dividing the mean μ_ddi_ by the standard deviation σ_ddi_ of dd_i_.

#### Equation (1)

For an exchange reaction ex_i_:

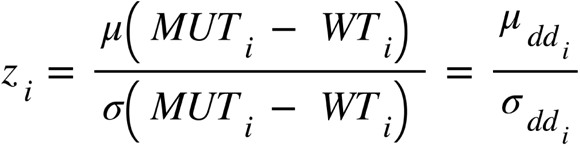

The z-score z_i_ is directional: a negative z-score indicates a decreased shift in the flux distributions from WT to MUT, and a positive z-score indicates an increased shift. A z-score close to 0 means that there is little difference between the distributions in the WT and MUT conditions. A z-score therefore represents the intensity and direction of one metabolite’s shift in a specific condition.

Here we propose the use of z-scores of all the exchange reactions in the network as a basis to rank all exchanged metabolites based on the intensity of the changes. Z-scores can also be used as-is or used with a threshold (see Discussion section). Since both increased and decreased metabolites are of potential interest, this ranking is based on the absolute values of the z-scores (Figure 2D). This reveals the metabolites whose import/export behaviour changes the most between the WT and MUT, relative to every other exchange metabolite in the network.

Furthermore, ranking the z-scores by absolute value provides insight via the comparison of the list of the top ranked metabolites between different scenarios, as ranks are relative to the entire list of exchange metabolites and not quantitative. In this paper, the rank calculated by SAMBA for each metabolite is referred to as SAMBARank.

### Applications to human metabolic profiling

In order to show the benefits of this methodology, we selected two complementary applications in human health. The goal was twofold: first to show the ability of SAMBA to retrieve known biomarkers, and then to show in an untargeted study that the newly introduced SAMBA ranking system is coherent with metabolomics data and indicate other potential metabolites of interest chemically related to the observed metabolites. First, the benefit of sampling for metabolic profile prediction is illustrated on a specific inborn error of metabolism (IEM) and associated biomarkers. Then, we use a mGWAS (metabolic genome wide association study) example to show how ranking these metabolite predictions can align with untargeted studies like metabolomics, and to recommend new potential metabolites of interest.

### Illustrating the benefits of sampling through the prediction of Xanthinuria type I biomarkers

Xanthinuria type I is a rare genetic disease caused by a mutation in the XDH gene [20], and is characterised by kidney stones (urolithiasis), urinary tract infections, and rarely kidney failure [21]. In patients with this disease, a decrease in urate and an increase in hypoxanthine has been observed (from OMIM [22]).

Here, we applied both FVA and sampling in order to compare the information which can be drawn from both techniques. Both the FVA bounds and the sampling distributions are displayed on the same plot for both of the expected biomarker metabolites (Figure 4). Expected biomarkers are defined as metabolites with observed significant changes in patients of the disease according to the original dataset. The flux simulations were run using Recon 2 [14], a human genome-scale metabolic network, by knocking out the XDH gene, which knocks out 7 reactions (see Table 1 in Methods). The flux values are reported on a log scale for clarity.

**Figure 4:**
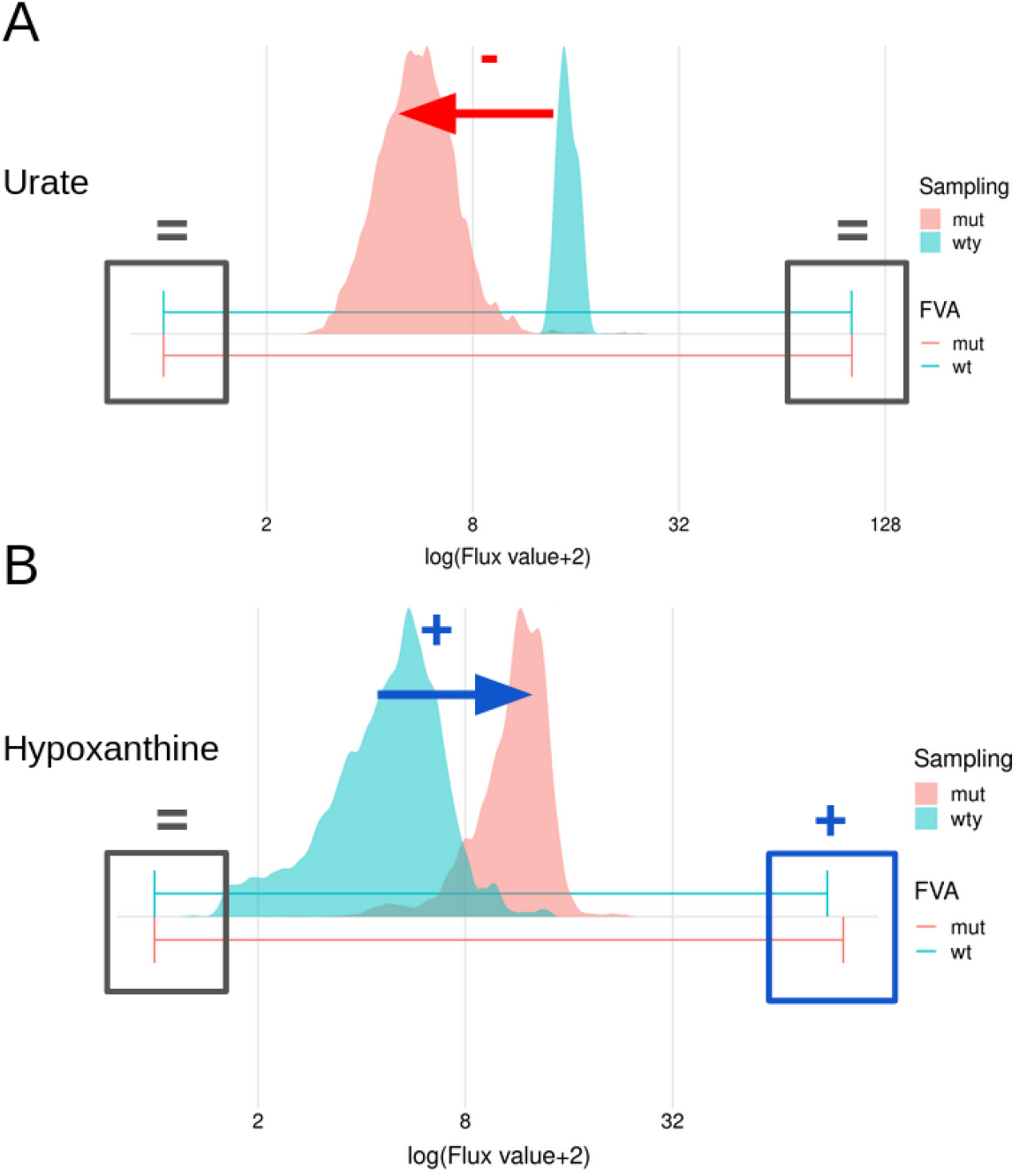
Flux bounds (FVA) and distributions (sampling) for urate and hypoxanthine in the WT state (light blue) and the MUT state (red). MUT here corresponds to the knock-out of the XDH gene. Highlighted in grey, red, and dark blue are the absences of shifts (=), decreases (-), and increases (+) respectively between WT and MUT.

**Table 1:** Genes and reactions knocked-out to simulate various conditions in metabolic models.

| Condition | Model | Gene KO | Reaction KO |
| --- | --- | --- | --- |
| Xanthinuria type I | Recon2 | XDH | XANDp XAO2x XAOx r0424 r0425 r0546 r0547 |
| SCD | Human1 (v1.14) | SCD, SCD5, FADS6 | MAR02281 MAR02282 MAR02284 MAR02286<br>MAR02287 MAR02292 MAR02293 MAR02294<br>MAR02295 MAR02296 MAR02288 MAR02289<br>MAR00144 MAR00146 MAR00147 MAR00148<br>MAR02126 MAR02128 |

For urate, the FVA bounds were the same in both conditions (Figure 4A), which is interpreted as a metabolite not considered as a biomarker in Shlomi *et al.* [13], and thus the FVA prediction does not agree with the observed decrease. On the other hand, the sampling distributions correctly show a decreasing shift from WT to MUT. For hypoxanthine, both methods are able to predict the expected increase in metabolite export, as shown in Figure 4B, via the shift in distributions for sampling and the change in upper bounds for FVA although this change is very small (10.3%) compared to the total feasible range.

Xanthinuria type I is one of many IEMs from an entire IEM - biomarker dataset which was curated in Sahoo *et al.* [23]. We used a subset of this dataset, used in Thiele *et al.* [14], to run our analyses by knocking out each gene responsible for each disease: we ran both FVA and sampling on the 49 IEMs for 54 metabolites on Recon 2. Heatmap figures containing the entire set of predictions for both FVA and sampling are included in supplementary data Figure S1, and overlaps are shown in Figure S2.

Overall, sampling not only complements FVA by providing new correct predictions, but also attributes more meaning to the scores of the predictions for each metabolite (see Figure S1). Sampling z-scores, as opposed to the binary increase/decrease indicators of FVA, can be used to rank, filter and gain insight on the intensity of changes. Indeed, once generated, distributions can be used in multiple ways, one of which is calculating z-scores and ranks as shown in this paper; in contrast FVA interval boundaries represent only the flux extremes instead of the general flux behaviour.

### Generating coherent recommendations and identifying novel metabolites of interest associated with SNPs using SAMBA

For this study, we chose a case example for the comparison of mGWAS data and SAMBA ranking system. In general, GWAS datasets are composed of traits associated with Single Nucleotide Polymorphisms (SNP), which are germline genetic substitutions of one nucleotide, present at a specific DNA position in at least 1% of the population. Specifically, mGWAS data consists of SNP-to-metabolic trait associations, such as single metabolite fold changes between non-SNP and SNP individuals, and ratios of two different metabolite levels, again compared between non-SNP and SNP individuals. Examples of these types of data are shown in Table 2. We used data from Suhre et al. [24], extracted SNPs associated with significant metabolites, and mapped them onto the Human 1 metabolic network.

**Table 2:** Example of mGWAS data: 3 significant metabolite ratios and 2 significant single metabolites. Beta’ represents the relative difference per copy of the minor allele (SNP) for the metabolic trait compared to the estimated mean of the non SNP population. The p-gain statistic quantifies the decrease in P value for the association with the ratio compared to the P values of the two separate corresponding metabolite concentrations.

| Ratio | beta' meta | P meta | p-gain meta |
| --- | --- | --- | --- |
| myristate (14:0) / myristoleate (14:1n5) | 0.124 | $2.9 \times 10^{-57}$ | $1.2 \times 10^{48}$ |
| myristate (14:0) / palmitoleate (16:1n7) | 0.131 | $1.4 \times 10^{-48}$ | $1.0 \times 10^{39}$ |
| margarate (17:0) / palmitoleate (16:1n7) | 0.157 | $2.1 \times 10^{-42}$ | $6.6 \times 10^{32}$ |
| margarate (17:0) | 0.06 | $4.9 \times 10^{-08}$ | 1 |
| myristoleate (14:1n5) | -0.075 | $3.3 \times 10^{-09}$ | 1 |

Among the 37 SNPs present in Supplementary Table 3 of Suhre et al., 17 were SNPs of 17 metabolic genes (one SNP per gene) present in the metabolic model Human 1 (version 1.10). The 20 other SNPs were impossible to simulate since they do not correspond to metabolic genes in Human 1. Human 1 was used to run sampling on 2 of these 17 SNPs: SNPs affecting the SCD gene and the ACADS gene. Human1 is one of the most recent and largest reconstructions of the human metabolic network, also showing that the method can scale to this larger network (13 024 reactions, 8 363 metabolites). The 15 remaining SNPs with corresponding genes in Human 1 were not analysed due to the manual curation needed to confirm genetic, enzymatic and metabolic matches.

We chose to focus on the SCD SNP specifically because i) the gene and reactions are present in the network, and ii) there are many measured metabolites present in the network, which is not the case for all of the SNPs, as some SNPs only have one or two significantly associated metabolites, or the associated metabolites do not exist in the network. It therefore serves as a good proof of concept application for the methodology. Furthermore, the selection of the correct genes to KO in the model for each SNP requires manual curation to make sure the GPRs (gene-protein-reaction) relationships correctly represent the enzyme and corresponding gene. Additionally, mapping the metabolite names from the study to model metabolites is a time consuming manual step. Results for SCD are shown in the main text, and those for the ACADS SNP are shown in Figure S4.

In contrast with IEM data, where mutations always result in an enzyme defect, an SNP might reduce enzyme activity (knock-down), enhance enzyme activity, or have an effect on a different gene. Some of the SNPs from the Suhre *et al.* study are well known to be associated with loss-of-function phenotypes such as enzyme deficiencies (*e.g.* the ACADS gene in ACADS-deficiency), and others have not been studied enough to confirm the effect of the SNP on gene function. As one example of an understudied SNP phenotype, the SCD gene (SNP rs603424 [25]) codes for the enzyme Stearoyl-CoA 9-desaturase, involved in fatty acid metabolism. The hypothesis is that the SNP mutation in the gene affects the corresponding enzyme negatively, which leads to no SCD enzyme activity, represented in the network by knocking-out the SCD gene and therefore blocking the corresponding reactions. This is suggested in Illig *et al.* [25] by drawing a parallel between known loss of function SNPs leading to severe disorders, and newly identified SNPs.

In Human 1, there are 19 reactions linked to the SCD gene, most of which involve the desaturation of stearoyl-CoA, palmitoyl-CoA and myristoyl-CoA into corresponding mono-unsaturated fatty acids. Following the GPRs relationships in the model, knocking out SCD only affects 4 reactions (due to the fact that SCD can be compensated by another gene). However, SCD also shares 14 GPRs with two other genes: SCD5 and FADS6, whose functions are not well described. We decided to knock out these extra 14 reactions in order to block the enzymatic function related to SCD completely.

The SCD SNP has two types of significantly associated metabolic traits: single metabolite changes, and ratios of two different metabolite concentrations. The single significant metabolites measured for the mGWAS study for this SNP are margarate, palmitoleate, myristoleate, stearate and 1-palmitoleoylglycerophosphocholine. These are the main “expected” metabolites, which will be compared with the SAMBA recommended metabolites.

SAMBA returned z-scores for the 1497 unblocked metabolite exchange reactions in Human 1. A metabolic profile this large is difficult to compare with the data from the mGWAS study as no raw data was included in the original study: only the significantly associated metabolites were reported. We also calculated the FVA bounds for each metabolite for the same metabolic condition as the sampling. Here, we compared the 5 significant metabolites reported in the mGWAS study with their simulated SAMBA metabolite ranks and FVA bounds to see the biggest effect this KO has on metabolite exports and imports.

Figure 5 shows the five metabolites identified in the mGWAS study along with the corresponding SAMBA ranks and the FVA predictions. Both in Figures 5 and 6, the metabolite(s) marked with “NA” in the SAMBARank column have no flux values because either they aren’t present as a metabolite in the network, don’t have an exchange reaction in the network, or have a blocked exchange reaction, meaning no flux can be carried through it in the current metabolic state.

**Figure 5:**
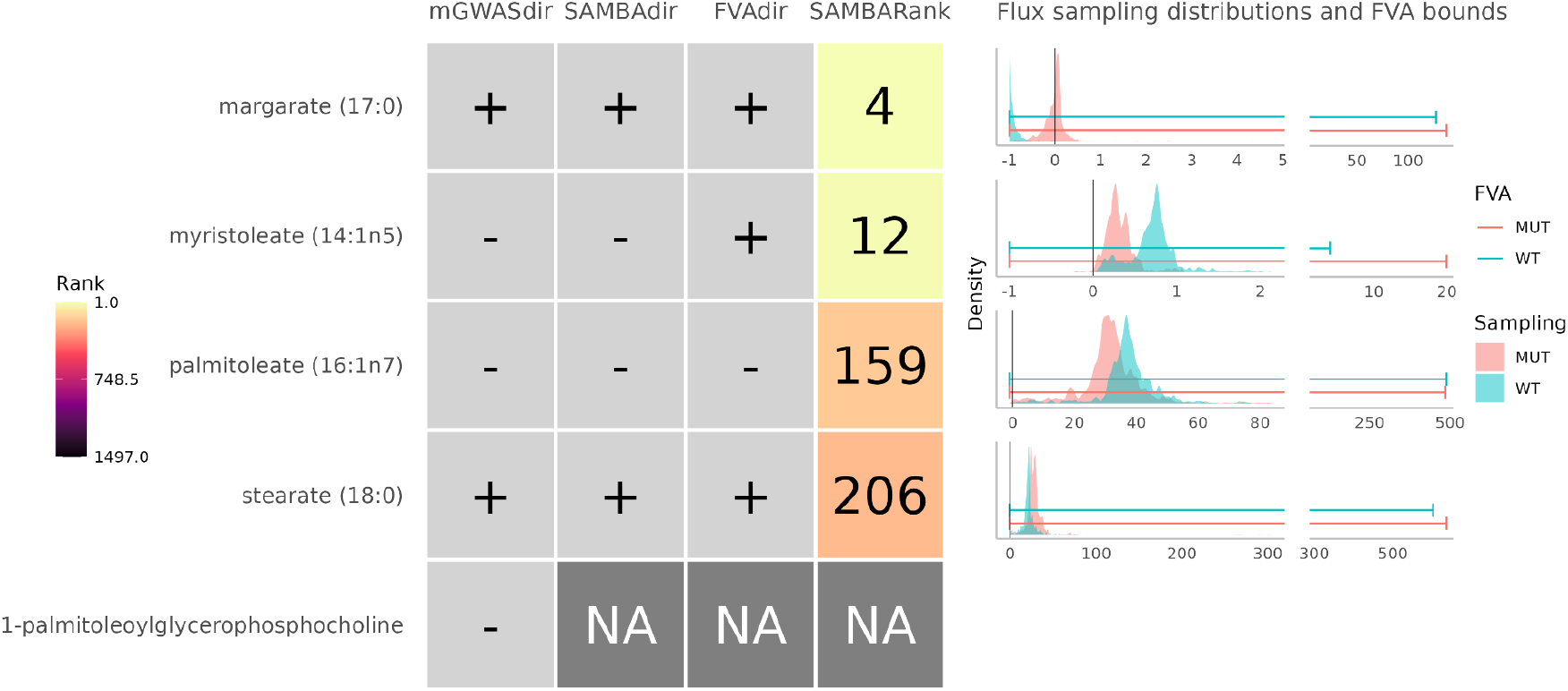
Observed and predicted changes for the five metabolites significantly associated with the rs603424 SNP. The first column shows the observed change directions from the mGWAS study. The second column shows the predicted change direction using SAMBA (SAMBAdir). The third column shows the predicted change direction using FVA (FVAdir). The fourth column shows the SAMBA predicted rank out of the 1497 metabolites in the network (SAMBARank). The NAs represent metabolites for which SAMBA was unable to predict fluxes for one of the following reasons: (i) the metabolite is not in the network, (ii) the metabolite is in the network but has no exchange reaction, or (iii) the metabolite’s exchange reaction can carry no flux (=blocked). Sampling distributions and FVA predicted bounds for each metabolite’s exchange reaction in WT and MUT are shown on the right.

**Figure 6:**
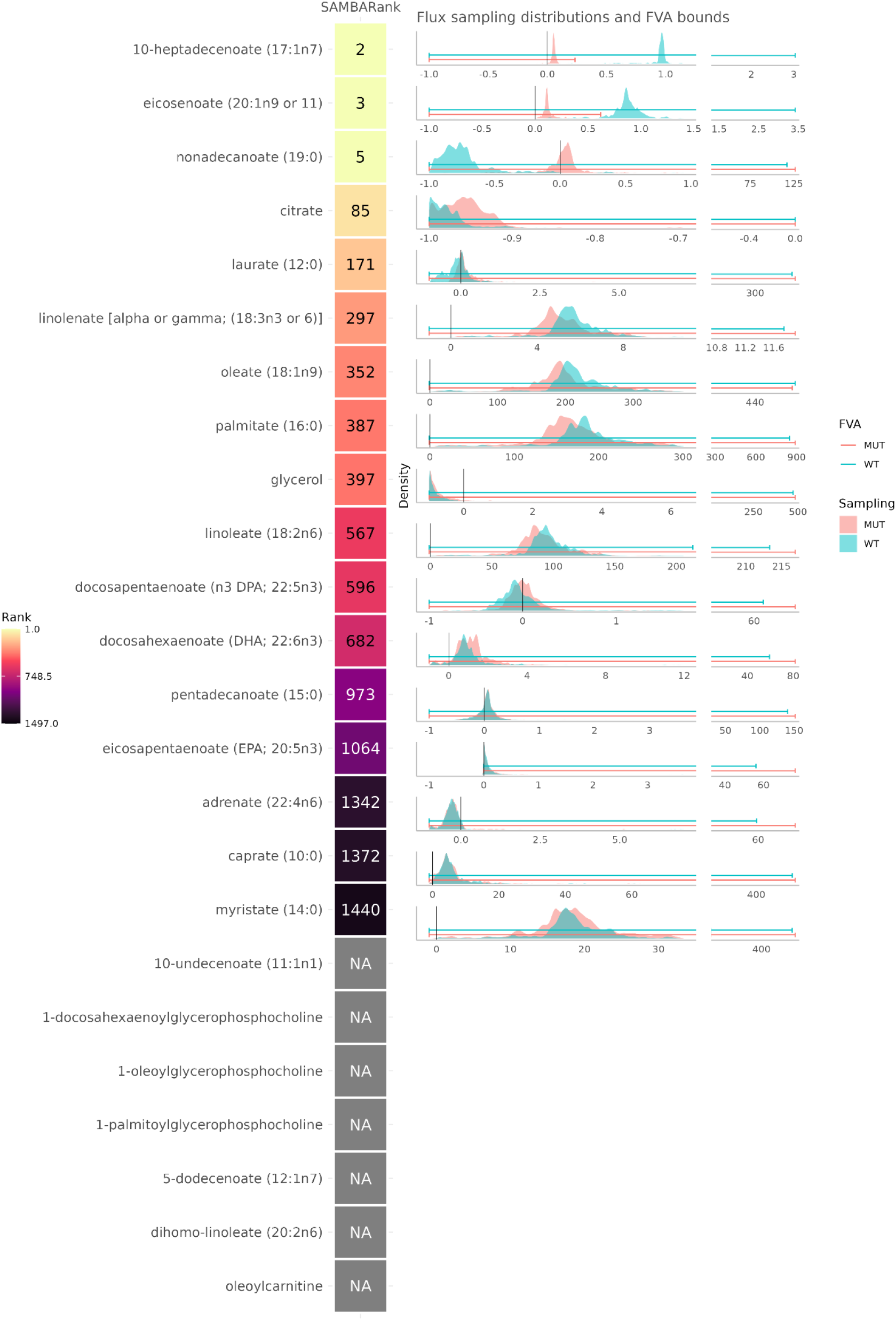
Predicted ranks for the metabolites present in a ratio significantly associated with the rs603424 SNP. The column shows the predicted rank out of the 1497 metabolites in the network. The NAs represent metabolites for which SAMBA was unable to predict fluxes for one of the following reasons: (i) the metabolite is not in the network, (ii) the metabolite is in the network but has no exchange reaction, or (iii) the metabolite’s exchange reaction can carry no flux (=blocked). Sampling distributions for each metabolite’s exchange reaction in WT and MUT are shown on the right.

Four out of the five expected metabolites are present with an exchange reaction in Human 1, and the SAMBA predicted change directions match the expected mGWAS experimental changes. The directions of change predicted by FVA are correct except for myristoleate, which was predicted to be increased instead of decreased using the FVA bounds. Their ranks are shown in the column SAMBARank and these ranks are to be compared with the total number of exchange metabolites present in Human 1, i.e. 1497. These four metabolites are in the top 13%, two of which are in the top 1%.

The significant metabolite ratios linked to SCD include many different combinations of pairs of metabolites. The assumption here is that at least one of the two metabolites involved in each ratio must change for the ratio to be significantly changed. Figure 6 shows the metabolites present in at least one ratio significantly associated with SCD and their associated predicted SAMBARanks.

The second most differentially abundant metabolite predicted by SAMBA for this condition is 10-heptadecenoate, which is present in at least one significant ratio in the mGWAS SCD dataset. In addition to this, there are 4 other highly ranked metabolites, all in the top 171 ranked metabolites out of 1497 (top 11%).

Furthermore, the top 10 most differentially changed metabolites associated with SCD predicted using SAMBA can be used to form a list of new metabolites of interest for this condition. By examining the chemical class of each predicted highly differentially abundant metabolite, we can gather information on a general type of metabolite affected by the KO.

Figure 7 shows the CHEBI ontology extracted using these top 10 metabolites. This hierarchical graph was made using BiNChE [26], which creates and enriches a subnetwork using a list of CHEBI IDs and the CHEBI ontology.

**Figure 7:**
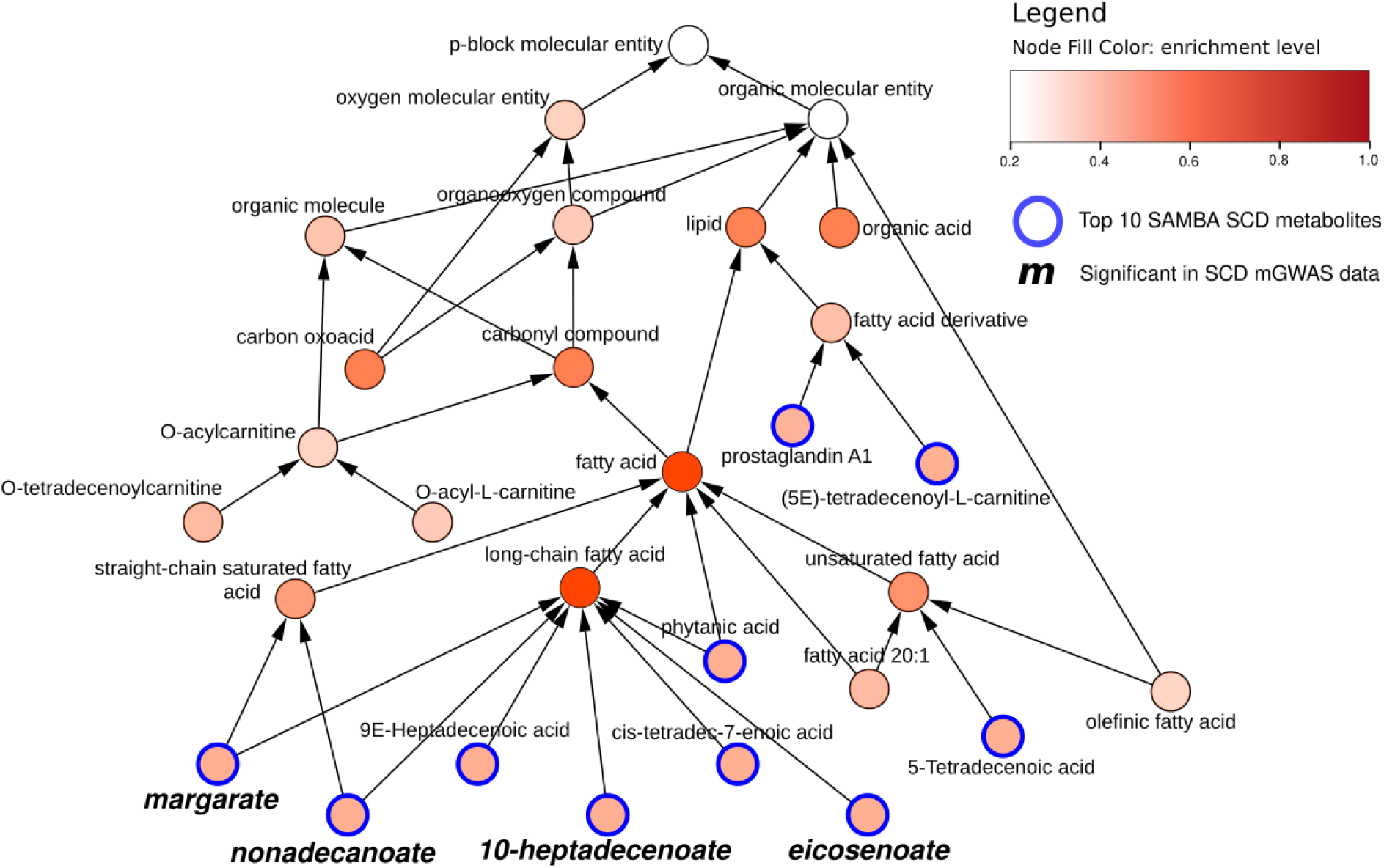
Hierarchical CHEBI graph of the top 10 metabolites predicted to be differentially abundant (outlined in blue) predicted by SAMBA for SCD, extracted using BiNChE. The node colour corresponds to the BiNChE enrichment level. The metabolites in bold & italic were significant in the mGWAS dataset for SCD.

All of the top 10 most changed metabolites are classed as lipids (outlined in blue in Figure 7), 8 of which are fatty acids, which is consistent with the functionality of the enzyme SCD. Indeed, the SNP rs603424 has been shown to be significantly associated with circulating phospholipid levels [27], as well as with low levels of palmitoleate [27]. SCD is a desaturase which leads to the formation of fatty acids, specifically monounsaturated fatty acids involved in membrane phospholipids [28].

Out of the top 10 metabolites, 4 were measured in the mGWAS study (margarate, 10-heptadecenoate, nonadecanoate, and eicosenoate), and they are all classified as saturated or long-chain fatty acids. This means that the other long-chain fatty acids could be potential metabolites of interest, such as 9-Heptadecenoic acid (rank 1) or cis-tetradec-7-enoic acid (rank 6), which weren’t measured in the original mGWAS study.

However, the ChEBI classification is limited by the annotation of each metabolite to the correct class. Upon manual inspection, both cis-tetradec-7-enoic acid and 5-tetradecenoic acid are C14:1 fatty acids, only differing by the position of the double bond, but they are classified separately in long-chain fatty acid and unsaturated fatty acid respectively. This indicates that 5-tetradecenoic acid could also be of interest for future studies. Furthermore, by looking at the chemical structures, 4 out of the top 10 are odd chain fatty acids which is interesting to highlight since they represent a very small percentage of the total human fatty acid plasma concentration [29].

Since BiNChE provides a view of the ChEBI ontology on a per-metabolite scale, using too many metabolites as input results in a large and difficult to read figure. Other methods can integrate more of the predicted metabolic profile (for example 50 metabolites).

As a step closer to using chemical structures as opposed to class annotations as well as using more of the metabolic profile, we ran a ChemRich [30] analysis using the top 50 metabolites predicted to be differentially abundant. It uses the chemical structure via SMILES, and the MeSH terms associated with PubChem IDs to highlight enriched chemical classes. Figure 8 represents the most enriched clusters from the top 50 metabolite set. The higher the -log(pvalue) (y axis), the more the group is enriched.

**Figure 8:**
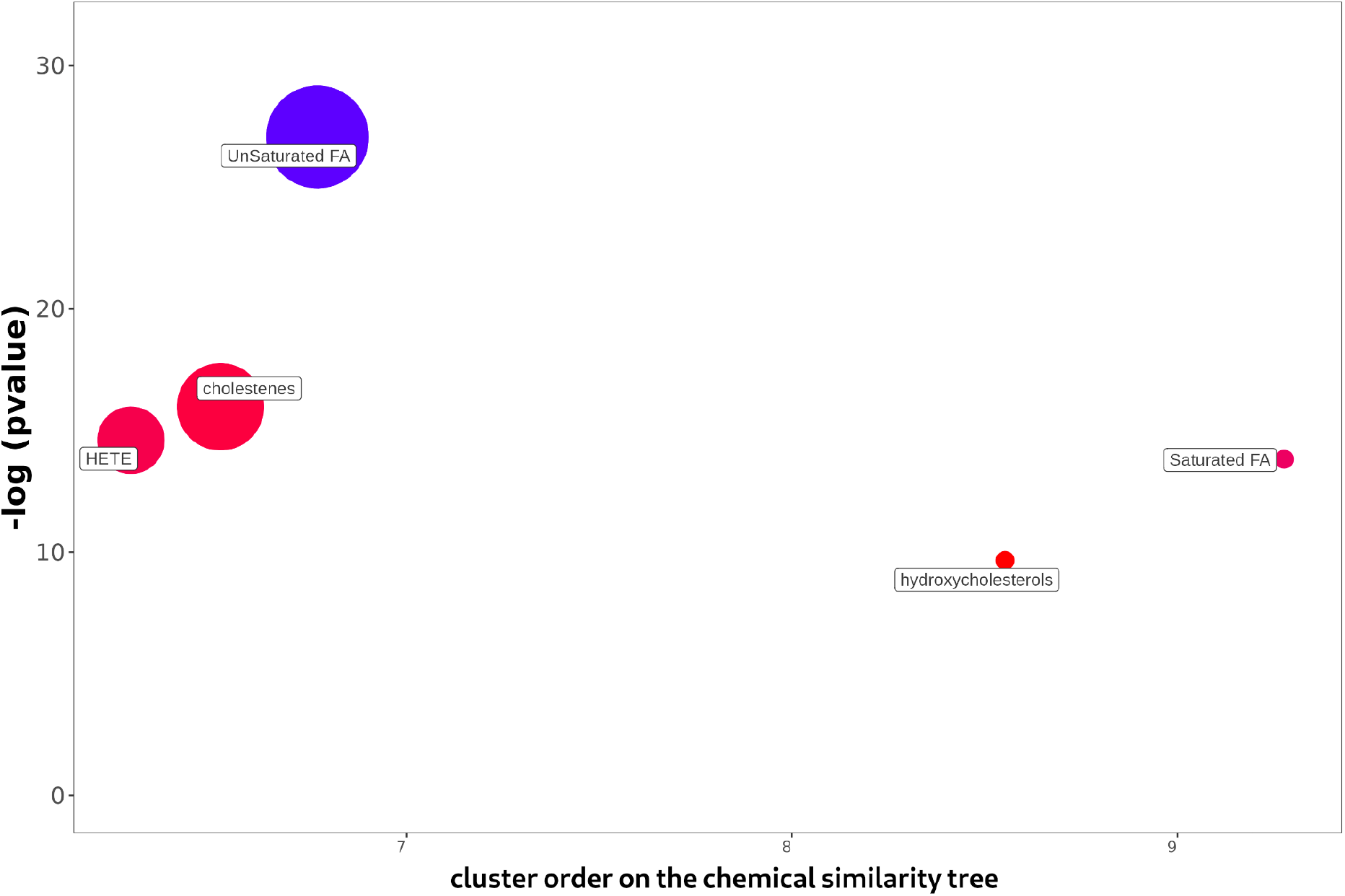
ChemRich enrichment of the top 50 most changed metabolites for SCD. The y-axis shows the most significantly altered clusters on the top. Each node reflects a significantly altered cluster of metabolites. Enrichment p-values are given by the Kolmogorov–Smirnov test. Node sizes represent the total number of metabolites in each cluster set. Cluster colours show the proportion of increased or decreased metabolites (red and blue respectively). The x axis represents a separation based on cluster order on the chemical similarity tree, and non-significant clusters are hidden.

The ChemRich plot also shows that both saturated and unsaturated fatty acids are significantly enriched by this dataset. Figure 8 also highlights some other groups such as HETE (Hydroxyeicosatetraenoic acids (which are oxylipins)), cholestenes, and hydroxycholesterols not detected using BiNChE. ChemRich serves as a complementary method to BiNChE for analysing predicted metabolic profiles, as highlighted in Table S2.

These are just two methods of enriching a metabolite set of interest. Moreover, as well as looking into specific metabolites as potential biomarkers, using SAMBA could direct future research in SCD towards the general families of unsaturated fatty acids and straight-chain saturated fatty acids, with wider panels of measurements for better coverage.

## Discussion

The results presented in this study show that by using metabolism-simulating methods like SAMBA, we can predict metabolic profiles. For instance, for the SCD case study, the metabolites reported as associated with the SNP were highly ranked, especially when considering the total number of exchange metabolites in the whole human network.

Metabolites belonging to the predicted list but not in the original metabolomics fingerprint may be of interest to improve metabolic profiling. In fact, as it was shown in Frainay *et al.* [31], metabolites may be overlooked during the whole metabolomics pipeline. This can be for instance due to pre-processing steps since most peak picking methods [32] will define an intensity threshold to keep only intense peaks and, as a consequence, may discard peaks of interest that fall just below the threshold.

The measurement of pure standards of metabolites is essential to obtain the highest level of confidence in metabolite identification (level 1 according to Metabolomics Standard Initiative [6]). Selecting which standards to measure is by itself a challenge, since samples can contain thousands of metabolites. Hence, SAMBA can be used by laboratories to select which standards to acquire in the context of the disease under study. More broadly, the top ranked list can also be used to identify families of metabolites to study as a whole, such as by extending the panel of measurable metabolites during a metabolomics experiment.

Although SAMBA is a predictive method, evaluating the predictions using traditional contingency tables, recall and precision is difficult due to the nature of metabolomics measurements and the available “truth” datasets. The model contains all known metabolites involved in metabolic reactions, but metabolomics methods are not able to detect and annotate all of them. This results in many cases where metabolites are predicted to be of interest while they are not detected by typical assays. In these cases, the predictions could be correct while being considered as a “false positive”. Instead of using “False Positive” to represent these predictions, we simply present the entire ranked prediction results in order to orient the user towards certain metabolites or metabolite classes. We then evaluate the method using true positive ranks and the list of the top most changed metabolites, some of which could be considered false positives, but could also be unmeasured metabolites.

SAMBA is based on ranking z-score absolute values, meaning that the metabolites whose exchange fluxes (and by extension concentrations) are more likely to change will be considered first. There are of course metabolites whose concentrations can change very little and have extreme consequences on the rest of the metabolism, such as via enzyme regulation, or if they are limiting substrates for example.

We demonstrated that sampling can add a layer of information to better improve metabolic profiling compared to FVA. Sampling provides a finer grained description of changes which helps order metabolites based on their likelihood to be affected by a perturbation. Compared with FVA, sampling is more computationally intensive (CPU and memory) but recent strategies are reducing this computational burden [33–35]. Nevertheless, sampling is currently more than feasible on large networks such as Human1.

The FVA method used in previous work [13] to compare intervals calculates the greatest change between the two pairs of upper bounds and the two pairs of lower bounds. This comparison of boundary shifts is not always representative of the underlying changes and can mislead the interpretation of the intensity of these changes. Using other methods such as comparing the means of boundaries assumes a uniform flux distribution within these bounds, which we have shown via sampling is rarely the case. Using the most frequent fluxes with sampling appears as a good approximation of the mix of metabolite exports that occurs in biofluids, but it should be noted that the most frequent flux value may not be the most frequently observed flux in reality. However, in some cases, the most frequently predicted flux value may not represent the biological reality of a cell, such as for cells in extreme conditions or fast-growing cancerous cells, for which fluxes might be more close to the extremes. To represent these extreme conditions in SAMBA, the initial parameters of the model could be adjusted (higher minimal production of biomass) to force the model to operate within extreme (boundary) optimums as opposed to more likely fluxes.

The boundary shifts evaluated by FVA are very sensitive to change, since a very low threshold (1e-6 tolerance and 0.01 factor) for change is used to report an increase or a decrease. Despite this, FVA is able to predict biomarkers, as shown in previous studies [13,14], when aiming to predict specific biomarkers. We progressed from the calculation of a score to the ranking of these scores since ranking the change intensities via sampling means that the most changed metabolites can be highlighted, while still keeping information on the other subtle metabolite changes. Contrary to the binary change/no change method of reporting FVA results, sampling ranks provide information on a wider scale by taking into account relative changes between metabolites.

In order to continue to highlight the full benefits of using sampling distributions instead of FVA boundary values, further research for other applications and more validation data are required. For instance, experimentally-measured *in vivo* fluxomics data [36] could be matched to simulated import/export rates. A database of every unique KO could be simulated and compiled as a repository for comparison with real data to determine which metabolic perturbations are most likely to cause the condition tested by the experiment.

This ranking system bypasses the issues that come with using flux values directly, and especially helps in choosing which metabolites to focus on first. The comparison of metabolic profile recommendations between different scenario simulations can be achieved by considering the top most changed metabolites and their ranks, as opposed to the raw flux values.

Z-scores prove to be useful in that they reflect an intensity of change similar to fold changes, and are weighted by the standard deviation of the distributions, which helps the z-scores to remain flexible given the variable nature of these distributions. Initially, instead of using a z-score to compare sampling distributions, more widely used statistical metrics were tested, such as Kullback-Leibler Divergence, Kolmogorov-Smirnov, and Wasserstein. However, they did not prove to be informative in our use case since they lead to p-values being too sensitive, resulting in extremely significant p-values for very similar distributions. In addition to this, these tests provide scoring metrics which are unable to quantify or describe the differences in the way a z-score can.

The goal of this study is to simulate whole-body metabolic markers using a generic genome-scale model. From a physiological point of view these models may seem to be somewhat over simplified in that a single metabolic system is represented. However the examples used in this study are genetic diseases, therefore they affect the genome of all of the cells in the body. While gene expression can depend on organs and tissue regions, the hypothesis here is that experimentally observed metabolic profiles are a combination of metabolite exports from all tissues connected to biofluids, which is why they can be equated to metabolic profiles predicted using a genome-scale network. However, the modulation of a tissue-specific biomarker may be predicted incorrectly if it is normally (biologically) compensated by other tissues, which could result in false positives. In those cases, tissue-specific networks could be useful for analysing diseases that are known to affect a certain tissue, such as glycogen storage diseases. These diseases are a collection of genetic metabolic disorders, and the enzymes affected by the mutations are specific to the liver and muscle [37]. By using transcriptomics data to create a liver-specific model, the accuracy of metabolic simulations could be increased. This can be done using various integration methods such as iMAT [38] or DEXOM [39]. However, choosing any given model and tissue-specific conditions must be done with care as it will have a major impact on the resulting metabolite ranks.

Furthermore, including the SAMBA approach in whole-body metabolic models [12] which combines the interactions of multiple human tissues is a potential path for future study. Since sampling algorithms are being continuously improved and iterated upon, and more CPU power is being added to computational clusters, running sampling on these larger models will become less of an issue. These models, with their different gene and reaction expressions per tissue, could reveal the different effects of genetic diseases or other metabolic disruptions on biofluid metabolites on a multi-tissular level.

Finally, while SAMBA was applied to KO scenarios in this paper, the method can be adapted to more complex constraints such as multiple gene KOs or even to simulate knock downs of reactions. This can be particularly useful in the context of toxicology or drug development, where these subtle metabolic disruptions can lead to reduced enzyme activity.

## Conclusion

Building upon constraint based modelling of metabolism through the use of random sampling of fluxes, we were able to predict large potential metabolic profiles and confirm measured metabolites both in targeted and untargeted assays. Ranking all metabolites becomes possible through the methodology’s comparison of flux distributions between healthy and disease states. Metabolites revealed by this method are of potential interest to broaden the panel of targets for future metabolomics experiments, and can be identified as understudied metabolites, helping to develop our understanding of metabolic mechanisms. Furthermore, the rank of a given metabolite can be compared between two different disruption scenarios, which provides information on the specificity of the disrupted metabolite to the scenario.

Although the methodology is designed to be used to predict external metabolite exchange fluxes, it can also be used to simulate the internal reaction fluxes, which can be useful for understanding internal metabolism along with external metabolites. Finally, simulated metabolic profiles can also be used to benchmark various analyses specific to metabolomics, such as pathway analysis, or other analyses which require lots of data like machine learning.

## Materials and Methods

### Metabolic models

In this study, Human 1 v1.14 [40], containing 13 024 reactions, was used to carry out mGWAS analyses. https://github.com/SysBioChalmers/Human-GEM Recon 2 [14], containing 7 440 reactions, was used to carry out IEM analyses. https://github.com/opencobra/COBRA.papers/tree/master/2013_Recon2

### WT and MUT states

When choosing which metabolic network to use, a decision must be made on whether to optimise the biomass reaction or not. This is done by providing the name of the reaction in the network (as they are not named uniformly between networks) as well as the fraction of biomass to optimise for, as SAMBA inputs. A list of genes or reactions to knock-out must also be provided to create the MUT state.

The WT state is created using the default network parameters (reaction bounds, biomass coefficients etc.). Then, the reaction(s) to be knocked out are forced to carry a non-zero flux. This is done by optimising for the reaction(s) to KO and then changing the minimum bound to 5% of the maximum flux value, or maximum bound to -5%, for forward and reverse reactions respectively. This avoids sampling and comparing two states where the fluxes are zero for the reactions of interest, and is also why each WT is specific to a MUT state.

The MUT state is created using the default network followed by setting the bounds for the reaction(s) to KO to zero.

### Model parameters

Sampling and FVA were run using the same parameters as in Shlomi *et al.* and Thiele *et al.* [13,14]: minimum fraction of optimum of the objective function set to 0, and all exchange reaction bounds set to [-1, 1000].

### Sampling method

Random sampling is done using Python code written for SAMBA, based on the cobrapy [41] Python package. The code uses the CPLEX 12.10 solver by default and uses the optGpsampler algorithm [42] to sample from the reaction flux solution space. optGpsampler begins with a warm-up phase to select starting points (by running a preliminary FVA on each reaction), followed by uniform sampling within this feasible solution space. Because each sample is selected from the solution space directly, there is no sample rejection since this would be extremely inefficient to do on genome-scale models. A thinning parameter of k (default k = 100) means that every k sample is saved and the rest is discarded in order to reduce intersample correlation. For large models such as Recon 2 and Human1, 100 000 samples with a thinning of 100 were used.

The number of samples can be changed, however, in order to sufficiently explore the solution space, a large number (at least 100 000) of samples must be used for larger networks such as Recon 2 or Human 1 which contain thousands of reactions. Determining a sufficient number of samples is discussed below in Sampling Convergence and supplementary data.

### Parallel processing

Sampling can be run on a local computer for smaller models, but it needs a certain amount of resources to run correctly. More specifically, the amount of RAM required increases with the size of the model, and more CPUs will help generate the samples faster.

For this study, the larger metabolic models (Recon2 and Human1) were sampled using a computer cluster using 16 cores and 128GB of RAM for each job. The cluster we used is the Genotoul computational cluster which has about 3000 cores / 600 threads, 36 Tera Byte memory (3TB on a SMP machine), Infiniband interconnection (QDR/FDR), parallel file system (GPFS).

### Sampling convergence

One of the main limitations of CBM is that it is impossible to fully describe such a large solution space. When using random sampling to explore the solution space, the number of samples to use must be provided, but choosing the ideal number for a given network is a challenge since by definition the structure of the solution space is unknown.

Therefore it is essential to know when to stop sampling: determining when the solution space has been sufficiently sampled. We ran convergence tests using various well-known sampling metrics: running means, traceplots, and shrink factor plots, to make sure that using 100 000 samples was enough for a network this large, for the goal of calculating z-scores on distributions. The results can be found in the supplementary data (Figure S3).

### Plots & libraries

The SAMBA pipeline is managed using Snakemake. Venn diagrams were made using the R library eulerr. Sampling distributions and mGWAS plot tables were made using ggplot2. BiNChE: the BiNChE plot was created by using a docker containing an old version of Firefox and Flash. The subnetwork was exported and then taken into Cytoscape [43] to change the layout, followed by Inkscape to edit the placement of labels.

### Code availability

The code for the SAMBA project is freely available at https://forgemia.inra.fr/metexplore/cbm/samba-project.

### Data

Using the mGWAS SNP dataset, genes were mapped to the Human 1 network using the annotated ENSG IDs. Then, using the model’s GPR relationships, reactions were automatically knocked out using SAMBA. In the case of SCD, the GPRs were manually checked. The SCD SNP only affects the SCD1 gene (known as SCD in the metabolic model), as SCD1 and SCD5 are two separate genes. SCD5 codes for the same enzymatic function as SCD1 but they are both expressed in different tissues, fat tissue for SCD1 and brain and pancreas for SCD5. However, Human1 is not tissue-specific and the reactions are not necessarily associated with the genes according to this tissue specificity, so in order to block the enzymatic function completely, both SCD1 and SCD5 were blocked.

Resulting SAMBA metabolites were manually mapped to the mGWAS significant metabolite names for SCD, with manual verification of metabolite synonyms as many lipids have multiple names and naming conventions.

## Acknowledgements

This work was supported by the French National Facility in Metabolomics & Fluxomics, MetaboHUB (11-INBS-0010). We would like to thank Jake Bundy for his participation in our discussions involving this work. The authors are grateful to the GenoToul bioinformatics platform (Toulouse, France) for hosting and providing their computer cluster, used for running all sampling runs.

## Supporting information

**File S1. Supplementary figures and tables.** Contains Figures S1, S2, S3, S4, and Tables S1 and S2.

**File S2. Full results for SCD metabolite SAMBA predictions.**

**File S3. Metabolite names and Human1 IDs for significant SCD metabolites.**

